# IL-32 is a stress-responsive node locked in a noncanonical NF-κB inflammatory loop during MASLD to HCC transition

**DOI:** 10.64898/2026.07.21.739717

**Authors:** Li Na Zhao, Philipp Kaldis, Jesper B. Andersen

**Affiliations:** Department of Health and Medical Sciences, Biotech Research and Innovation Centre (BRIC), University of Copenhagen, Copenhagen, Denmark; Department of Clinical Sciences, Clinical Research Centre (CRC), Lund University, Malmö, Sweden

**Keywords:** IL-32, NF-κB, lipid metabolism, MASLD, HCC, DGAT2

## Abstract

**Background:** Interleukin-32 (IL-32) presents a long-standing paradox in liver disease, with markedly elevated expression in hepatocellular carcinoma (HCC) yet a protective role against hepatic steatosis. The absence of a canonical receptor or defined secretory pathway has obscured its biological function. This study aimed to resolve this paradox by delineating the regulatory mechanisms that govern IL-32 activity during hepatocarcinogenesis.

**Methods:** We analyzed two in-house prospective cohorts, including a MASLD cohort and a MASLD-associated HCC cohort, integrating matched transcriptomic and metabolomic data. Targeted lipidomics and multi-omics analyses were combined with single-cell and spatial transcriptomics. Key findings were validated using functional assays and gene perturbation models.

**Results:** We identified a disease stage-specific transcriptional switch in which noncanonical NF-κB signaling (NFKB2/RELB) replaces canonical NF-κB as the primary activator of IL-32, forming an auto-amplifying inflammatory loop. This switch is enabled by FOXO1, which acts as a pioneer factor to maintain chromatin accessibility at IL32 and NF-κB loci. Functionally, IL-32 is coupled to lipid metabolism through DGAT2; however, this axis becomes uncoupled in HCC, where DGAT2 loss rewires NF-κB/ERK signaling without recapitulating global metabolic remodeling, thereby sensitizing cells to inflammatory activation.

**Conclusions:** These findings resolve the functional paradox of IL-32 by revealing a multi-layered regulatory network that reprograms its activity during liver disease progression, and define IL-32 as a context-dependent integrator of metabolic and inflammatory signaling, whose regulatory network is rewired during hepatocarcinogenesis to promote a sustained pro-inflammatory state.

**Highlights:**

- Noncanonical NF-κB (NFKB2/RELB) drives a self-amplifying IL-32 loop in HCC.
- FOXO1 licenses this switch by maintaining chromatin accessibility at IL32 and NF-κB loci.
- IL-32 shifts from a metabolic regulator in MASLD to an inflammatory driver in HCC.

## INTRODUCTION

Many interleukins drive progression from metabolic dysfunction-associated steatotic liver disease (MASLD) to hepatocellular carcinoma (HCC) by sustaining chronic inflammation, yet the mechanistic roles of most remain incompletely defined (1–3). Among these, IL-32 stands out: it shares no sequence homology with other cytokine families and lacks a conventional receptor (4–6). Initially characterized in immune cells, recent studies have revealed that IL-32 directly regulates triglyceride (TAG) metabolism and fibrogenic signaling in human hepatocytes, positioning IL-32 as a candidate molecular integrator of metabolic and inflammatory circuits (7).

IL-32 is expressed as multiple isoforms (including IL-32α, β, γ, δ, and ε), each showing distinct potencies and biological specialization (8,9). Among these isoforms, IL-32γ induces high levels of the pro-inflammatory molecules (TNF, IL-6, IL-1β, and CXCL8) (10). Each isoform can activate NF-κB but differ markedly in their capacity to modulate apoptosis, antiviral immunity, and crosstalk with pattern-recognition receptors (PRRs) (11).

MASLD represents a global health crisis, with many patients advancing to steatohepatitis (MASH) with severe fibrosis, and ultimately HCC (12). This progression is driven by the predominant pathological convergence of lipid overload and chronic, unresolved hepatic inflammation (13,14), as well as context-dependent dysregulation of epigenetic and transcriptional networks (15,16).

Chronic inflammation is a well-established driver of hepatocarcinogenesis (17), with the NF-κB transcription factor family occupying the central role in this process (18). Canonical NF-κB signaling (RELA/p50) has garnered the vast majority of attention as the primary mediator of pro-inflammatory responses, whereas the non-canonical pathway (RELB/p52) remains comparatively underexplored in the context of liver cancer (19–22). IL-32 is a pro-inflammatory cytokine known to activate NF-κB, yet its role in liver disease presents a striking paradox. IL-32 is elevated in MASLD patients and further augmented in MASLD-HCC (23), suggesting a pathogenic function. However, genetic ablation of IL-32 promotes hepatic steatosis (24), indicating a protective role in metabolic liver disease. This contradictory involvement, protective in steatosis but induced in cancer, underscores the urgent need to define how IL-32 shifts from a metabolic regulator to an inflammatory driver during hepatocarcinogenesis.

To resolve this paradox, we harnessed two in-house clinical cohorts (MASLD and MASLD-HCC) with matched hepatic and serum transcriptomic/metabolomic data, complemented by scRNA-seq, spatial transcriptomics, genetic perturbation, and epigenomic profiling. We identified a disease stage-specific transcriptional switch in which non-canonical NF-κB (NFKB2/RELB) supersedes the canonical pathway as the dominant driver of *IL-32* in HCC, establishing a self-reinforcing inflammatory circuit. We further demonstrate that this switch is licensed by FOXO1 through chromatin decompaction, a pioneer factor function previously unrecognized in this context.

## METHOD

### Patient cohorts

We collected two human cohorts spanning the MASLD progression spectrum: a MASLD cohort (**n=114**, of which 90 have matched RNA-seq and liver/serum metabolomic profiles) and a paired MASLD-HCC cohort comprising tumor and adjacent non-tumor tissues (**n=47** pairs).

### Ethics statement

The MASLD study protocol conforms to the ethical guidelines of the 1975 Declaration of Helsinki and was approved by Etikprövningsmyndigheten (Dnr 2024-07917-01) in Sweden. MASLD-HCC was performed following individual patient consent, local institutional review board approvals (IORG0003254; IRB00003888), and assessed by the Committee on Health and Research Ethics for the Capital Region of Denmark for use of archival material ( no. H-4-2016-FSP, 17029679). All patient datasets were anonymized.

### MASLD metabolomics

Metabolomic profiling of the cohort samples was performed using our established protocols as previously described (13). Briefly, sample processing and data acquisition were conducted on LC-MS/MS platforms. The raw data were processed through a standardized pipeline for peak picking, alignment, and normalization against internal standards and quality control samples. Compound identification was performed by matching against authentic chemical standards and curated metabolite databases. The processed metabolomic datasets from the MASLD cohort was subsequently processed for downstream integrative analysis. Data frames for liver, and plasma metabolites, alongside gene expression data were created by selecting only the samples (n=90) present across all three data modalities.

### Targeted lipidomic profiling of paired MASLD-HCC tissues

To characterize the metabolic remodeling specific to hepatocellular carcinoma, we profiled a paired tumor and non-tumor liver tissue cohort using targeted lipidomics (performed by OWL, Spain).

### RNA-sequencing

Transcriptome sequencing was performed by Biomarker Technologies (BMK) GmbH. Total RNA was extracted from liver biopsies using TRIzol reagent. RNA quantity and purity were assessed using a NanoDrop spectrophotometer, and RNA quality was evaluated using a Labchip GX system. cDNA libraries were prepared using the Hieff NGS Ultima Dual-mode mRNA Library Prep Kit for Illumina (Yeasen) according to the manufacturer’s protocol. Paired-end sequencing (150 bp) was conducted on a NovaSeq 6000 platform. Raw sequencing data were quality-checked with FastQC (v0.11.9). Reads were aligned to the human reference genome GRCh38 (Ensembl release 77) using STAR (v2.7.9a), and gene expression levels were quantified as transcripts per million (TPM) using RSEM (v1.3.1). For downstream analyses, TPM values were log₂-transformed [log₂(TPM + 1)] and mean-centered.

### Statistical analysis and bioinformatic methods

Detailed descriptions of all statistical and bioinformatic analyses are provided in the Supplementary Information. Statistical analyses were performed using methods appropriate for each dataset and experimental design, and the specific statistical test used for each analysis is indicated in the corresponding figure legend.

## RESULTS

### IL-32 is elevated in hepatic steatosis and involves TAG remodeling

To investigate how IL-32 relates to hepatic lipid accumulation and coordinated inflammation (**Fig. 1A**), we performed RNA-seq on a MASLD cohort (n=114 samples) stratified by steatosis severity (S0-S2; n=70). Hepatic *IL-32* expression progressively increased with steatosis severity (**Fig. 1B**). Untargeted metabolomic profiling of transcriptome-matched liver tissues revealed that hepatic IL-32 levels were specifically associated with the accumulation of medium-chain triglycerides (TAGs) in the liver, and this association strengthened proportionally with the degree of TAG polyunsaturation (**Fig. 1C**). This observation is restricted to the intrahepatic lipid pool as no association was found either by lipidomics evaluating plasma TAG species or by clinical biochemistry measuring circulating TAG levels (**Fig. S1a-b**). This indicates that IL-32 expression (dictated by the isoforms IL-32β and IL-32γ) is more tightly coupled to local hepatic lipid remodeling rather than the systemic lipid burden (**Fig. S1c**).

**Figure 1.**
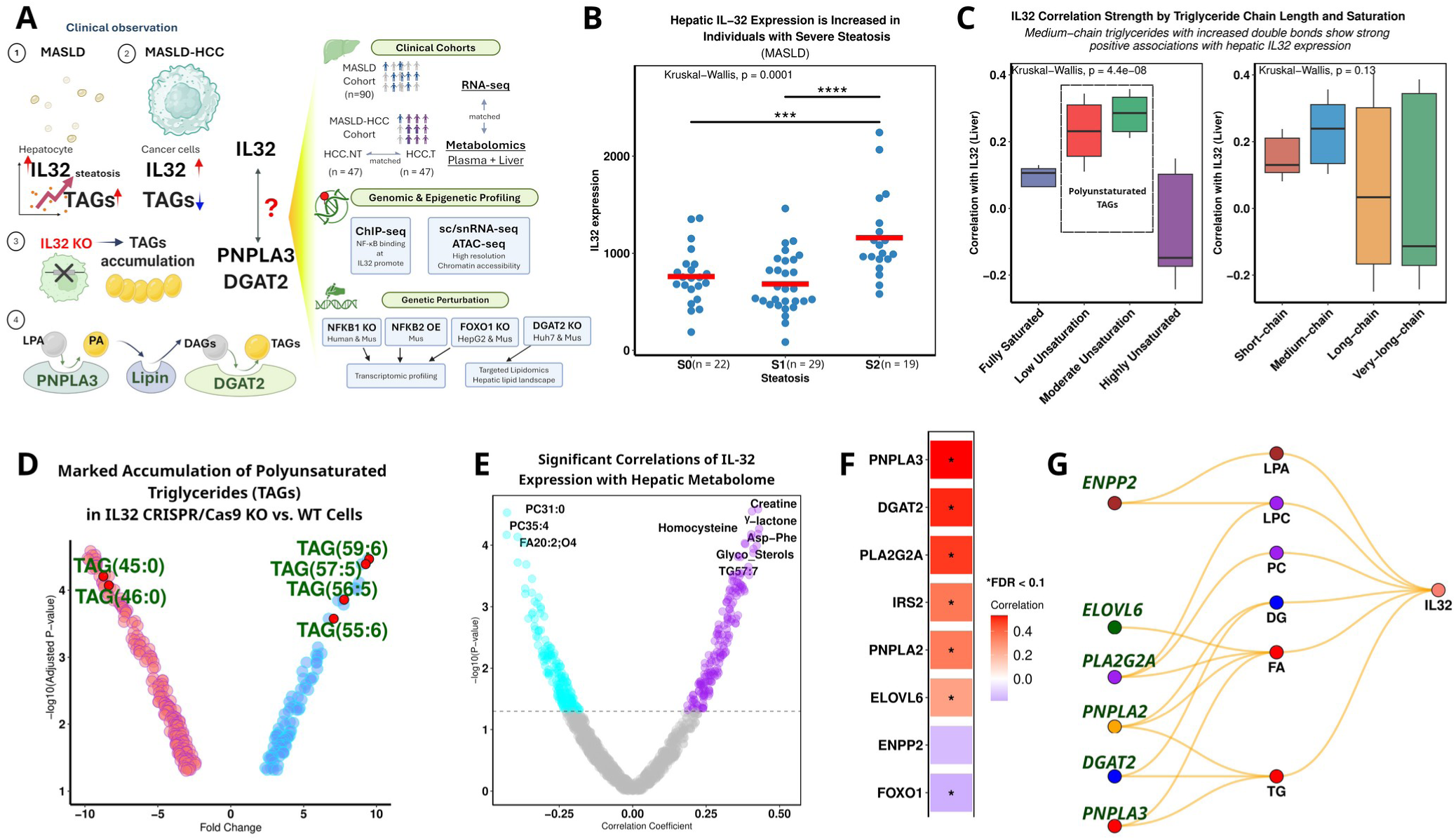
IL-32 is a steatosis-associated hub gene linked to TAG remodeling. (A) Study key questions and design. All human hepatic transcriptomic, lipidomic, and metabolomic data presented in panels B, C, and E-G were derived from the MASLD patient cohort. (B) Hepatic *IL32* expression across degrees of steatosis. (C) Hepatic *IL32* correlations with hepatic TAG species by unsaturation and chain length. Left panel: correlation strength of IL-32 with TAGs grouped by degree of unsaturation. Right panel: correlation of *IL32* with triglycerides by chain length. (D) Volcano plot showing differential lipid species between *IL32* CRISPR/Cas9 knockout (KO) and wildtype (WT) cells. Polyunsaturated TAGs significantly altered in KO cells are highlighted in red. (E) Hepatic *IL32* correlations with hepatic metabolites. Each point represents a metabolite, with the x-axis showing the correlation coefficient and the y-axis showing statistical significance (-log10 P-value). (F) Top correlations between hepatic *IL32* expression and metabolically relevant enzymes. (G) *IL32*-centered enzyme-metabolite network.

### Multi-omics analysis identifies IL-32 as a central regulator of metabolic rewiring in MASLD

To characterize metabolic pathways linked to IL-32 control in MASLD, we performed integrated transcriptomic and metabolomic analyses using pairwise correlation matrices to generate gene-metabolite association networks (**Fig. 1D-E**). This analysis positioned IL-32 at the core of the enzymatic regulation, in which metabolites serve as intermediates that link key hepatic enzymes controlling lipid processes to IL-32 (**Fig. 1F-G**). This IL-32-centered network highlights two main axes of regulation that are directly supported by significant positive correlations shown in human liver tissues. These include TAG remodeling that predominantly is associated with PNPLA3 and DGAT2 as well as oxidative lipid signaling, which is routed through PLA2G2A (**Fig. 1G**, **S1d**). Moreover, genetic knockout of *IL32* is shown to significantly reduce DGAT2 expression levels (**Fig. S1e**). Beyond metabolic enzymes, IL-32 (particularly the IL-32β and IL-32γ isoforms; **Fig. S1d**) is significantly correlated with the C-C motif chemokine ligand 20 (CCL20) (**Fig. S1f**), a critical proinflammatory and immune regulatory molecule (25). Taken together, these data raise the question of whether IL-32 coordinates a linked network between lipid metabolism and hepatic immune signaling.

### IL-32 upregulation is linked to lipid remodeling in MASLD-associated HCC

Since IL-32 has been shown to be central in the TAG remodeling, we next investigated how *IL32* transcription is regulated during progression from MASLD to MASLD-associated HCC. Tumor transcriptomes obtained from patients with MASLD-HCC were compared to matched non-tumoral (HCC.NT; n=47 paired samples) tissues, revealing a substantial remodeling of the IL-32 regulatory landscape in HCC (**Fig. 2A**). This finding was validated in transcriptomes obtained from TCGA-LIHC (n=377). Besides, the metabolic enzymes shown to be significantly associated with the *IL32* transcription, we additionally found NF-κB family members showing highly elevated expression levels in HCCs (**Fig. 2B**), which suggests that the IL-32 regulatory network is rewired during MASLD-to-HCC progression. Although IL-32 is markedly elevated in tumors at both the transcript and protein levels (an upregulation predominantly driven by the IL-32γ isoform), its associated metabolic network is disrupted in HCC (**Fig. 2B-C**). Specifically, we found in HCC a direct absence of the positive association between IL-32 and DGAT2 levels, with some targets (such as *PNPLA3*) showing inverse expression at both the mRNA and protein levels (**Fig. 2A-B**).

**Figure 2.**
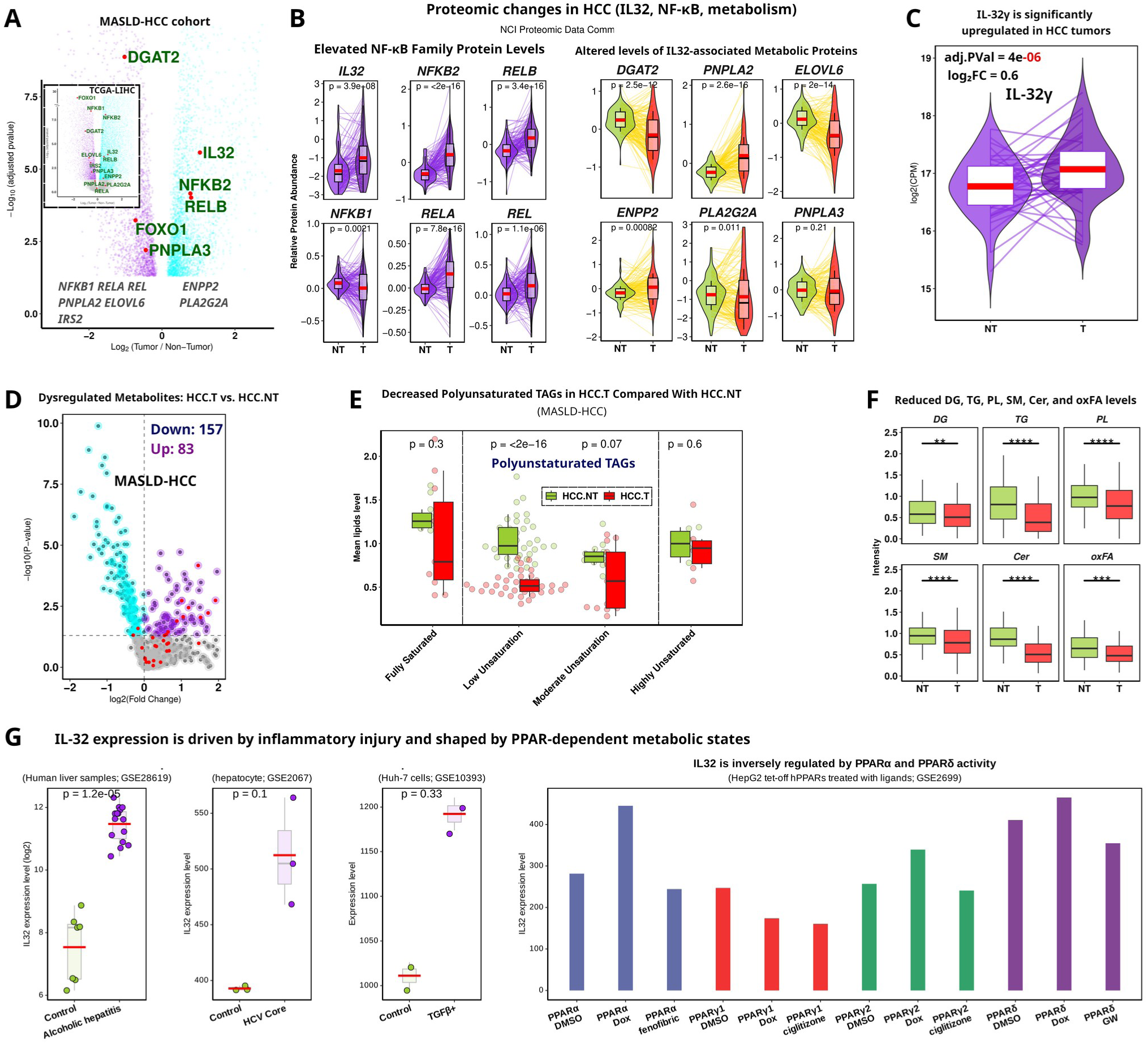
Dysregulation of IL-32 and NF-κB signaling and lipid remodeling in liver cancer. (A) Volcano plots showing downregulated *PNPLA3* and *DGAT2* and upregulated *IL32* and NF-κB noncanonical pathways (*NFKB2*, *RELB*) in the MASLD-HCC cohort comparing non-tumor to tumor samples. Top corner panels display corresponding expression patterns in TCGA-LIHC for validation. (B) Changes in protein abundance of IL-32, NF-κB family, and *IL32*-associated metabolic proteins in liver cancer using data from the NCI Proteomic Data Commons (PDC) CPTAC-HCC cohort. (C) *IL32*γ expression in HCC tumor tissue compared with non-tumor liver. (D) Volcano plot showing downregulated and upregulated metabolites in HCC tumors versus non-tumor tissue (HCC.T vs. HCC.NT). Red color indicates fatty acids. (E) Comparison of TAG levels across double bond numbers in HCC tumors versus non-tumor liver. (F) Reduced lipid species in tumor hepatocytes. (G) IL32 increases in alcoholic hepatitis (GSE28619), HCV core protein stimulation in hepatocytes (GSE2067), and TGF-β-treated Huh7 cells (GSE10393), and exhibits subtype-specific responsiveness to PPARα, PPARδ, and PPARγ activation, indicating sensitivity to both inflammatory and metabolic regulatory signals (GSE2699).

Targeted lipidomics revealed that IL-32 upregulation is accompanied by extensive hepatic lipid remodeling in HCC. Tumors displayed 83 lipids significantly elevated, including essential and non-essential amino acids, N-acyl ethanolamines, and beta-hydroxy acids, suggesting selective accumulation of bioactive and signaling lipids (**Fig. 2D**, **S2a**). Conversely, 157 lipid species were found significantly decreased, including polyunsaturated TAGs, diacylglycerides, phospholipids, and sphingolipids (such as ceramides and monohexosylceramides), reflecting a widespread depletion of storage and structural lipids (**Fig. 2E-F, S2b**). Matched transcriptomes confirmed a corresponding and significant downregulation of key enzymes, including the terminal TAG synthase (*DGAT2*), fatty acid oxidation (FAO) enzymes (*CPT1A, CPT2, ACADM, ACADL*), and the lipase *MGLL* (**Fig. S2c**). The concurrent depletion of lipid species, including major membrane phospholipids (such as PCs and PEs), and reduced expression of their regulatory enzymes that are accompanied by a significant augmented IL-32 expression, indicates a significant metabolic rewiring from lipid anabolism and storage towards lipid species that promote proliferation, signaling, and energy homeostasis in HCC cells (**Fig. S2d**).

IL32 expression is consistently elevated in inflammatory and injury-associated contexts, including alcoholic hepatitis, HCV core protein stimulation, and TGFβ-treated hepatocytes (**Fig. 2G**). To further assess whether IL32 is also responsive to metabolic regulatory signals, we examined datasets involving activation of peroxisome proliferator-activated receptor (PPAR) subtypes, key transcriptional regulators of hepatic lipid and glucose metabolism (26). Activation of PPARα and PPARδ suppressed IL-32, whereas their pharmacological suppression (e.g., by doxorubicin) elevated IL-32; conversely, PPARγ1 consistently reduced IL-32 regardless of ligand treatment, while PPARγ2 showed an intermediate response (**Fig. 2G**).

### Noncanonical NF-κB signaling takes control of IL-32 expression in HCC

To investigate upstream drivers of the IL-32 increase, we next examined the regulation of the NF-κB/Rel transcription family that previously has been implicated in *IL32* transcriptional control (27). ChIP-seq analysis shows a direct mechanistic connection to RelA, and NFKB2/p52 occupancy at the *IL32* promoter region, which is overlapping with the CAGE-defined transcription start sites and ATAC-seq accessible chromatin peaks, indicating that *IL32* regulatory elements are actively engaged by NF-κB (**Fig. 3A**). The noncanonical NF-κB pathway (NFKB2/p52) functions as a heterodimer together with RelB, suggesting that the observed NFKB2 binding represents a recruitment of the RelB-p52 complex at the *IL32* locus. Together with RelA, a core component of the canonical NF-κB pathway, these data indicate that both the canonical and noncanonical NF-κB signaling regulate *IL32* transcription.

**Figure 3.**
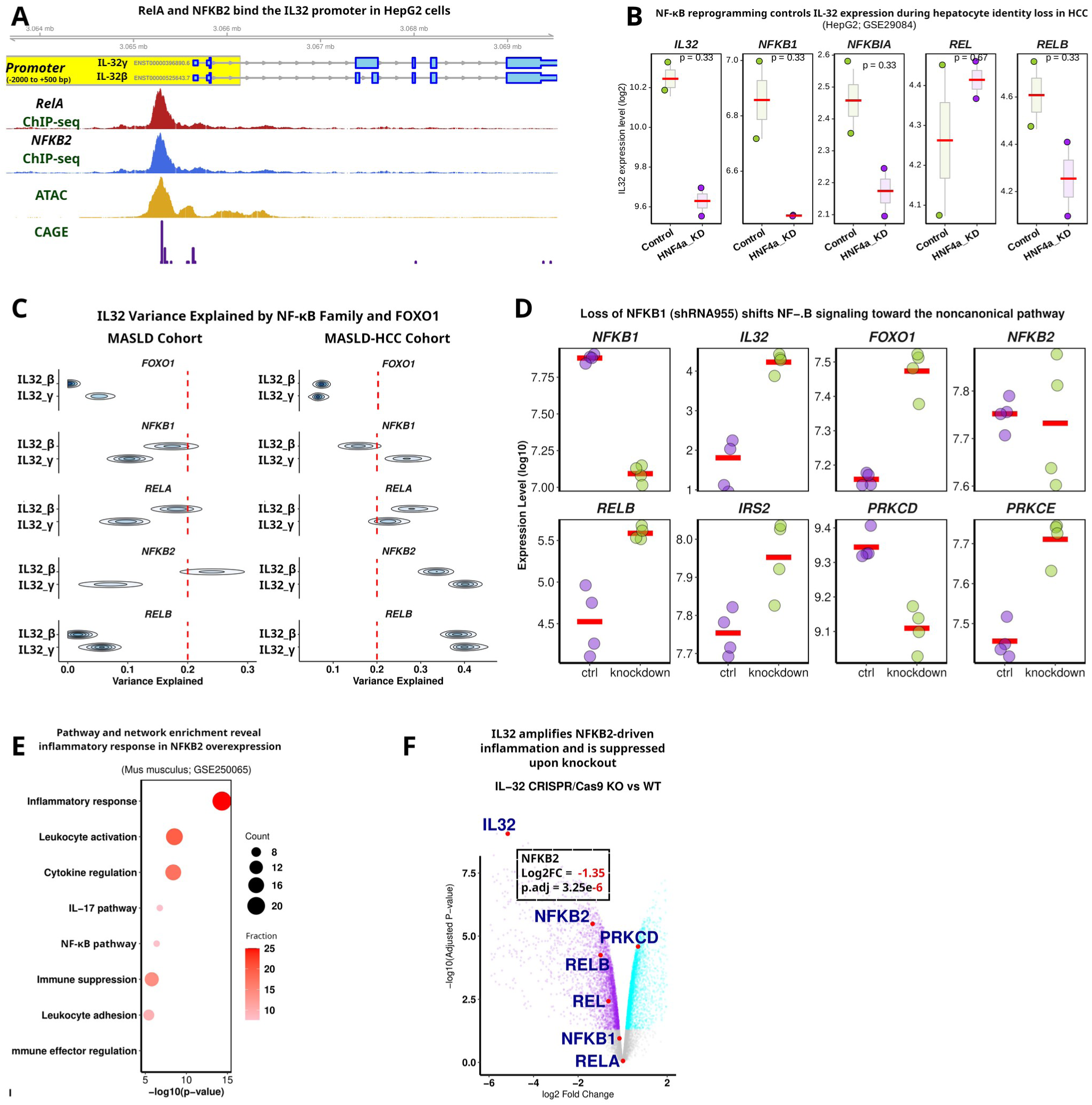
NF-κB regulation of IL-32 and pathway rewiring upon perturbation. (A) RelA and NFKB2 binding at the IL-32 canonical transcript in HepG2 cells. Gviz tracks display gene structure, promoter annotation, chromatin accessibility (ATAC-seq), and transcription start site activity (CAGE) across the IL-32 locus. ChIP-seq signal tracks for RelA and NFKB2 reveal promoter-proximal peaks at the canonical IL-32 transcript. Chromatin accessibility is shown using HepG2 ATAC-seq (ENCFF285FQS), while transcription initiation is indicated by CAGE signal (ENCFF205BRW). (B) Expression of selected NF-κB-associated genes following HNF4A knockdown in HepG2 cells (GSE29084). Statistical significance was assessed using the Wilcoxon test. (C) Density plot showing the proportion of *IL32* isoform expression variance explained by individual NF-κB family members and *FOXO1* in MASLD and MASLD-HCC cohorts. (D) Log-transformed expression levels of key inflammatory, metabolic, and signaling genes in control versus *NFKB1* shRNA (shRNA954; shRNA955) conditions. (E) Pathway enrichment analysis following *NFKB2* overexpression. Top 100 significantly differentially expressed genes were analyzed using Metascape. Results are based on the Mus musculus dataset from GSE250065. (F) Volcano plot showing differential NF-κB family gene expression in *IL32* CRISPR/Cas9 knockout (KO) versus wild-type (WT) samples.

Consistently, perturbation of hepatocyte identity signaling further supports this regulatory dependency (**Fig. 3B**): HNF4α knockdown resulted in coordinated downregulation of NFKB1, NFKB1A, RELB, and IL32, accompanied by compensatory upregulation of REL (c-Rel), indicating that IL32 expression depends on a balanced NF-κB network state rather than a single linear NF-κB branch.

To quantitatively assess the drivers of *IL32* transcription in MASLD and progressed HCC, we fitted a partial least square (PLS) regression model and assessed the predictive performance using 10-fold cross-validation. This analysis revealed that, in both the MASLD and MASLD-HCC patient cohorts, the noncanonical NF-κB components (NFKB2 and RELB) explain a substantially greater proportion of the variance in IL-32 expression (up to 40% of the IL-32 expression variance), than what can be explained by the canonical NF-κB factors (NFKB1 and RELA) (**Fig. 3C**).

Moreover, to examine these regulatory relationships, we analyzed IL-32 expression in *NFKB1* knockout human MCF7 cells (28). Loss of *NFKB1* was associated with increased expression of IL-32, FOXO1, NFKB2, and RELB, including reduced levels of IRS2, PRKCD, and PRKCE, which are consistent with a shift to an active noncanonical NF-κB pathway (**Fig. 3D**, **S3a**). Under these conditions, the IL-32 expression level correlates significantly with NFKB2/RELB and FOXO1, suggesting that IL-32 in a context-dependent manner may be controlled both by the canonical and noncanonical NF-κB signaling. These findings support a model, in which regulatory inputs to IL-32 shift during progression from MASLD to MASLD-associated HCC. Notably, the compensatory noncanonical NF-κB activation and FOXO1 upregulation are not observed in a murine model (**Fig. S3b)**, indicating that the rewiring of NF-κB signaling upon *Nfkb1* loss likely is tissue- and species-specific.

### A bidirectional regulatory loop amplifies inflammation in HCC

Functional studies have established *NFKB2* as a direct driver of inflammation (29). Moreover, ATAC-seq of *NFKB2* knockout cells did not reveal any changes in chromatin accessibility at the *IL32* and *DGAT2* loci (**Fig. S3c**). In contrast, murine *Nfkb2* overexpression showed a robust activation of inflammatory programs, with strong enrichment of pathways including leukocyte activation, cytokine production, IL-17 signaling, and NF-κB signaling itself (**Fig. 3E**, **s3d**). These results indicate that elevated NFKB2 in HCC directly drives a pro-inflammatory state, acting as a potential driver of IL-32 upregulation and functioning as an amplifier of immune activation during tumor progression. In contrast, *Nfkb2* overexpression causes marginal effects on the expression of key metabolic genes, including *Irs1, Irs2, Dgat2, Pnpla2, Elovl6, Enpp2,* and *Pnpla3* (**Fig. S3e**).

Importantly, a genetically engineered IL32 knockout model shows a significant reduction in both *NFKB2* and *REL* expression levels (**Fig. 3F**). In contrast, canonical NF-κB members (NFKB1 and RELA) remain statistically unchanged, suggesting a bidirectional feedback loop in which IL-32 reinforces noncanonical NF-κB activity.

### FOXO1 is a pioneer transcriptional factor regulating the *IL32-DGAT2* axis

Genetically engineered FOXO1 knockout HepG2 cells showed a significant reduction in both NFKB1 and REL expression levels (**Fig. S4a**), supporting a role for FOXO1 in modulating NF-κB signaling (30). Given the established role of FOXO1 regulating the hepatic metabolism and its predicted binding site within the *IL32* promoter (**Fig. 4A**), we next investigated whether FOXO1 regulates *IL32* transcription. Specifically, we examined whether FOXO1 influences NF-κB recruitment to the *IL32* promoter and how this regulatory axis links inflammatory signaling with the lipid dysregulation observed in MASLD-HCC.

**Figure 4.**
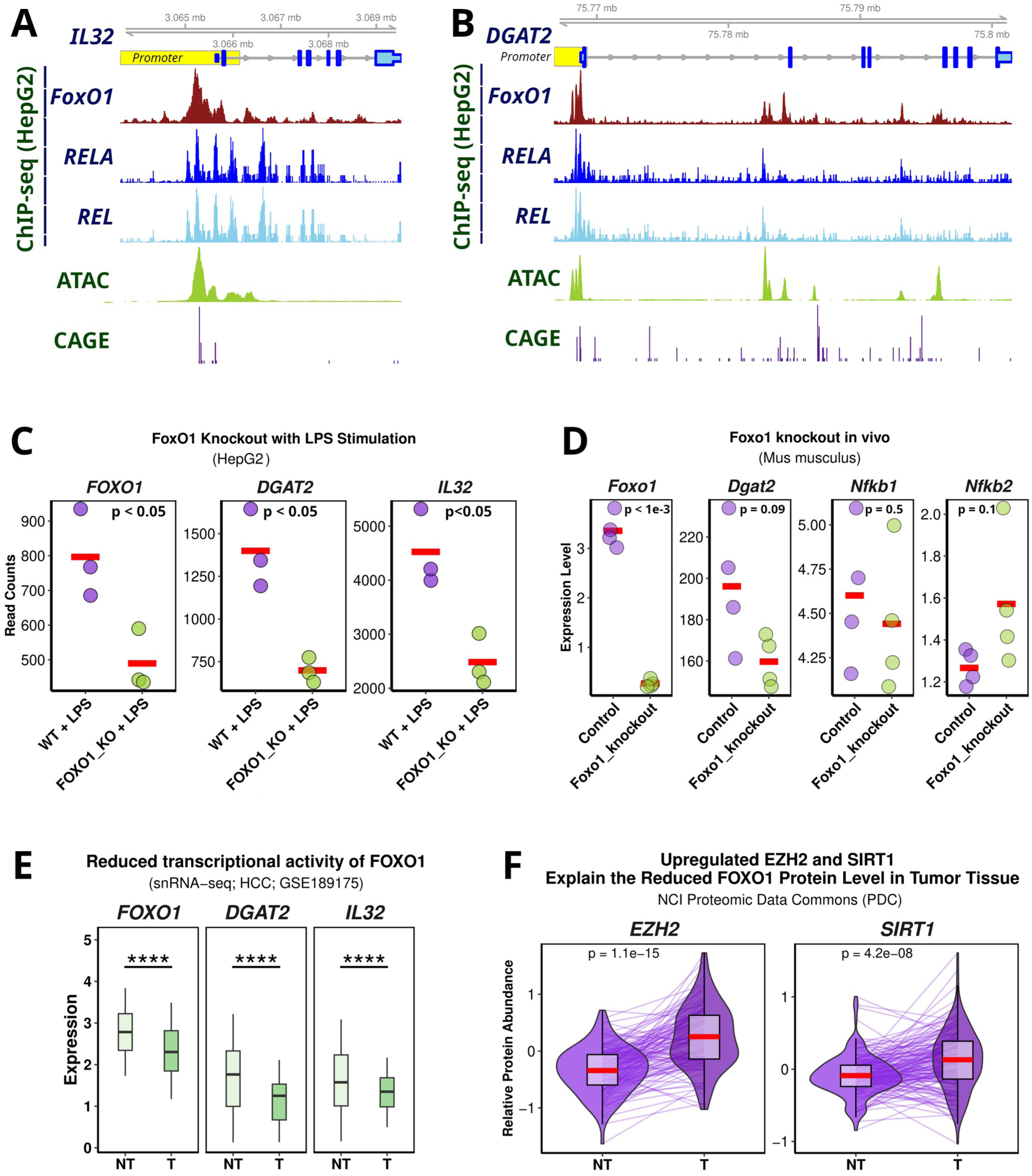
FOXO1-dependent regulation of IL-32 and DGAT2 in human hepatocytes and HCC. (A) Genomic view of the *DGAT2* promoter region showing co-localized chromatin and transcription factor activity in HepG2 cells. FOXO1, RELA, and REL signals are plotted together with HepG2 ATAC-seq accessibility and CAGE transcription initiation signal. Peaks from all datasets overlap at the DGAT2 promoter. (B) Genomic view of the *IL32* promoter region showing co-localized chromatin and transcription factor activity in HepG2 cells. (C) *FoxO1* knockout in LPS-stimulated HepG2 hepatocytes (GSE293003) results in downregulation of *IL32* and *DGAT2*. (D) *Dgat2* expression in hepatocyte-specific *FoxO1* knockout mice (GSE90754). (E) snRNA-seq analysis of snap-frozen archived liver biopsies from paired tumor and adjacent non-tumor tissues of 3 patients with fatty liver disease and hepatocellular carcinoma (HCC; n=3 pairs; GSE189175) shows that *FOXO1* expression is reduced in HCC, accompanied by decreased DGAT2 and IL-32. (F) Paired proteomic measurements from the NCI Proteomic Data Commons (PDC) showing EZH2 and SIRT1 protein abundance in HCC tumor samples versus matched adjacent normal tissue, n=277 paired samples.

In HepG2 cells, ChIP-seq against FOXO1 showed a direct binding at the promoters of both *IL32* and *DGAT2* (**Fig. 4A-B**). These regions significantly overlap with strong ATAC-seq peaks, indicating open chromatin accessibility, and coincide at the transcription start sites with positive CAGE signals, which is consistent with active transcription of the *IL32* and *DGAT2* genes. Importantly, each promoter regions are also co-occupied by the NF-κB subunits RELA and REL, suggesting that FOXO1 and NF-κB may jointly regulate the transcription of *IL32* and *DGAT2*.

To determine the functional consequences of this regulation, we assessed the transcriptional outcomes of FoxO1 depletion in liver cancer cells. In HepG2 cells, *FOXO1* knockout under lipopolysaccharide (LPS)-stimulation resulted in the significant downregulation of both DGAT2 and IL-32 expression (**Fig. 4C**). Furthermore, this data was corroborated in mice by hepatocyte-specific *FoxO1* knockout, which showed a significant downregulation of *Dgat2* (**Fig. 4D**). *In mente*, the absence of a murine *Il32* ortholog precludes its assessment in this model. Moreover, knockout of *FOXO1* in HepG2 was found to suppress the canonical NF-κB pathway, with marked downregulation of NFKB1 and REL (**Fig. S4a**). Consistently, the murine *FoxO1* knockout model showed reduced *Nfkb1* expression, alongside a trend toward increased expression of *Nfkb2* and *Relb*, suggesting a compensatory shift toward the noncanonical NF-κB pathway (**Fig. S4b**). Together, these findings demonstrate that FOXO1 is required not only for the expression of its direct transcriptional targets but also for maintaining canonical NF-κB signaling, positioning it as a regulator of this inflammatory cascade.

### Upstream EZH2 and SIRT1 drive FOXO1 silencing in HCC

In HCC, FOXO1 expression is reduced while AKT-mediated phosphorylation of FOXO1 remains unchanged, indicating that *FOXO1* silencing occurs independently of upstream AKT signaling (**Fig. 2A, Fig. S4c**). This regulation is evident at the single-nucleus RNA level, where FOXO1, DGAT2, and IL-32 are simultaneously downregulated in isolated malignant hepatocytes (**Fig. 4E**).

Next, we sought to determine the regulators responsible for *FOXO1* silencing. Among key regulators, only Hepatocyte Nuclear Factor 4-Alpha (HNF4A) and EZH2 (the histone methyltransferase, Enhancer of Zeste Homolog 2) were upregulated, whereas SIRT1 (the deacetylase, Sirtuin 1) was markedly downregulated in tumors compared to matched surrounding liver tissues (**Fig. S4d**). However, further analysis by ChIP-seq revealed only a weak HNF4A occupancy at the *FOXO1* promoter and no detectable CAGE transcriptional initiation (**Fig. S4e**), suggesting limited direct influence by HNF4A on regulating *FOXO1*. Similarly, HNF4A was also shown to only weakly bind the *DGAT2* promoter, which is in contrast with its observed functional impact on DGAT2, implying a context-dependent or indirect regulation (**Fig. S4f-g**). Contrary, EZH2 has been shown to directly bind and transcriptionally silence *FOXO1* via a PRC2-dependent H3K27me3 deposition (31). Besides, EZH2 expression inversely correlates with FOXO1 (**Fig. 4F**, **S4c**). Besides, EZH2 expression inversely correlates with FOXO1 (**Fig. 4F**, **S4c**). Moreover, studies have shown that SIRT1 overexpression reduces FOXO1 protein levels, and SIRT1 is known to mediate FOXO1 post-translational stability in addition to deacetylation-dependent functional regulation (32–35).

Taken together, the upregulation in HCC tumors of EZH2 and SIRT1 proteins provide a coherent mechanism for the reduced FOXO1 expression observed at both mRNA and protein levels (**Fig. 4F**). EZH2 mediates *FOXO1* promoter-level transcriptional silencing, whereas SIRT1 is known to mediate FOXO1 protein abundance, establishing a dual repressive axis driving FOXO1-depletion in HCC that operates independently of AKT phosphorylation.

### FOXO1 maintains chromatin accessibility at NF-κB target promoters

In HepG2 cells, *FOXO1* knockout significantly reduces NFKB1 and REL expression levels (**Fig. S4a**). Furthermore, FOXO1 co-occupies the *DGAT2* and *IL32* promoters with NF-κB subunits (RELA and REL) (**Fig. 4A**). To investigate the direct impact of FOXO1 on chromatin accessibility at high resolution, we leveraged a murine *FoxO1* knockout T cell model compared to its wild type cells (36), allowing for a comprehensive integration of ChIP-, ATAC-, and RNA-seq data. Utilizing these datasets, we showed a substantial *FoxO1* binding at promoters of *Dgat2*, *Nfkb1*, *Rela*, and *Relb* with corresponding open chromatin in wild type cells (**Fig. 5A-C**). Strikingly, *FoxO1* knockout significantly reduced ATAC signals at these promoter regions consistent with a loss of the accessibility and a reduced expression of all NF-κB family members (**Fig. 5C-D**). These analyses demonstrate that FOXO1 is required to establish and maintain an open chromatin configuration at the promoters of key metabolic and inflammatory genes, whereas *FOXO1* loss results in chromatin compaction and consequently downregulation of the NF-κB-controlled inflammatory gene-network (**Fig. 5E**).

**Figure 5.**
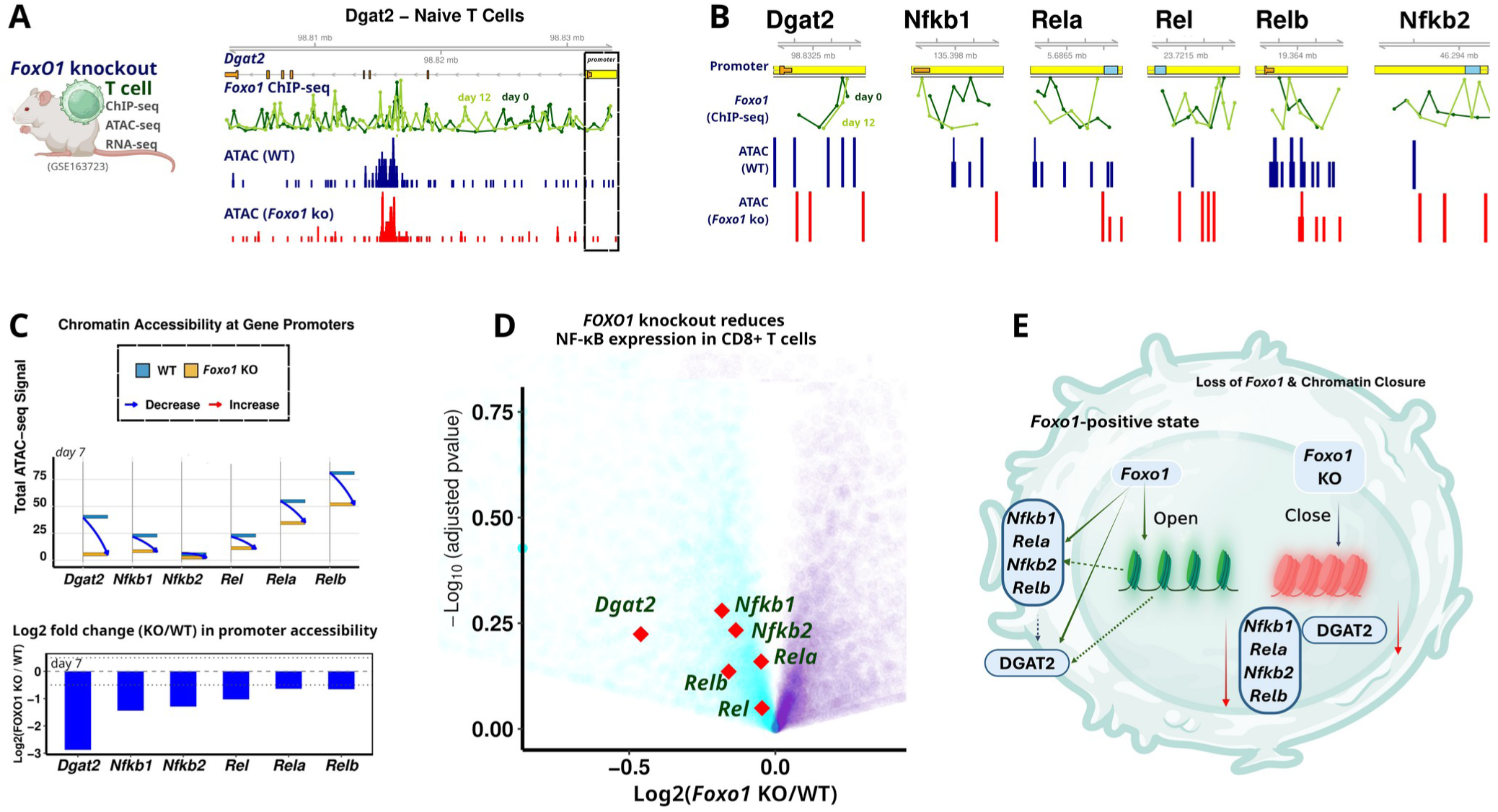
FOXO1 maintains chromatin accessibility for metabolic and NF-κB genes, and its loss enables inflammatory rewiring. (A) ATAC-seq (GSE163723) of the *Dgat2* promoter shows strong chromatin accessibility in wildtype cells, which is reduced in *Foxo1* knockout (KO) cells. Foxo1 ChIP-seq demonstrates binding at the *Dgat2* promoter overlapping regions losing accessibility in knockout. (B) ATAC-seq shows accessible chromatin at *Nfkb1, Rela, Nfkb2*, and *Relb* promoters in wildtype cells, but reduced in *Foxo1* knockout cells. Foxo1 ChIP-seq confirms binding overlapping these promoters. (C) Chromatin accessibility across gene promoters in wild type and *Foxo1* knockout cells (GSE163723). Upper panel: total ATAC-seq signal. Lower panel: log₂(Foxo1-KO/WT) ratio illustrating regions losing or gaining accessibility. (D) Volcano plots of NF-κB pathway gene expression in *Foxo1* knockout CD8⁺ T cells compared to wild type. (E) Model: FOXO1 sustains chromatin accessibility for metabolic and NF-κB genes.

### *DGAT2* loss is insufficient for metabolic remodeling but rewires NF-κB/ERK signaling in IL-32-high HCC

DGAT2 expression is tightly correlated with IL-32 and functionally linked to TAG production (Fig. 1F). Consistent with its role in lipid storage, DGAT2 expression is dynamically regulated by multiple metabolic inputs, including increased expression upon high-fat diet (HFD) feeding and Lipin1β overexpression, and responded to altered fatty acid desaturation (SCD1 deficiency), indicating that DGAT2 expression reflects both lipid availability and lipid composition states (**Fig. 6A**). Given that IL-32 upregulation in HCC is associated with coordinated metabolic remodeling, we hypothesized that DGAT2 loss may function downstream of IL-32 signaling to modulate lipid-dependent sensitivity to inflammatory cues (**Fig. 6B**). To test this, we analyzed Dgat2 knockdown in mice with established MASLD, associated with increased activity of NF-κB and ERK-responsive transcriptional regulators (Egr1 and Junb) alongside downregulation of Lpin2, which control lipid metabolism and are involved in inflammation (**Fig. 6C**), consistent with a shift toward a pro-inflammatory and metabolically dysregulated state.

**Figure 6.**
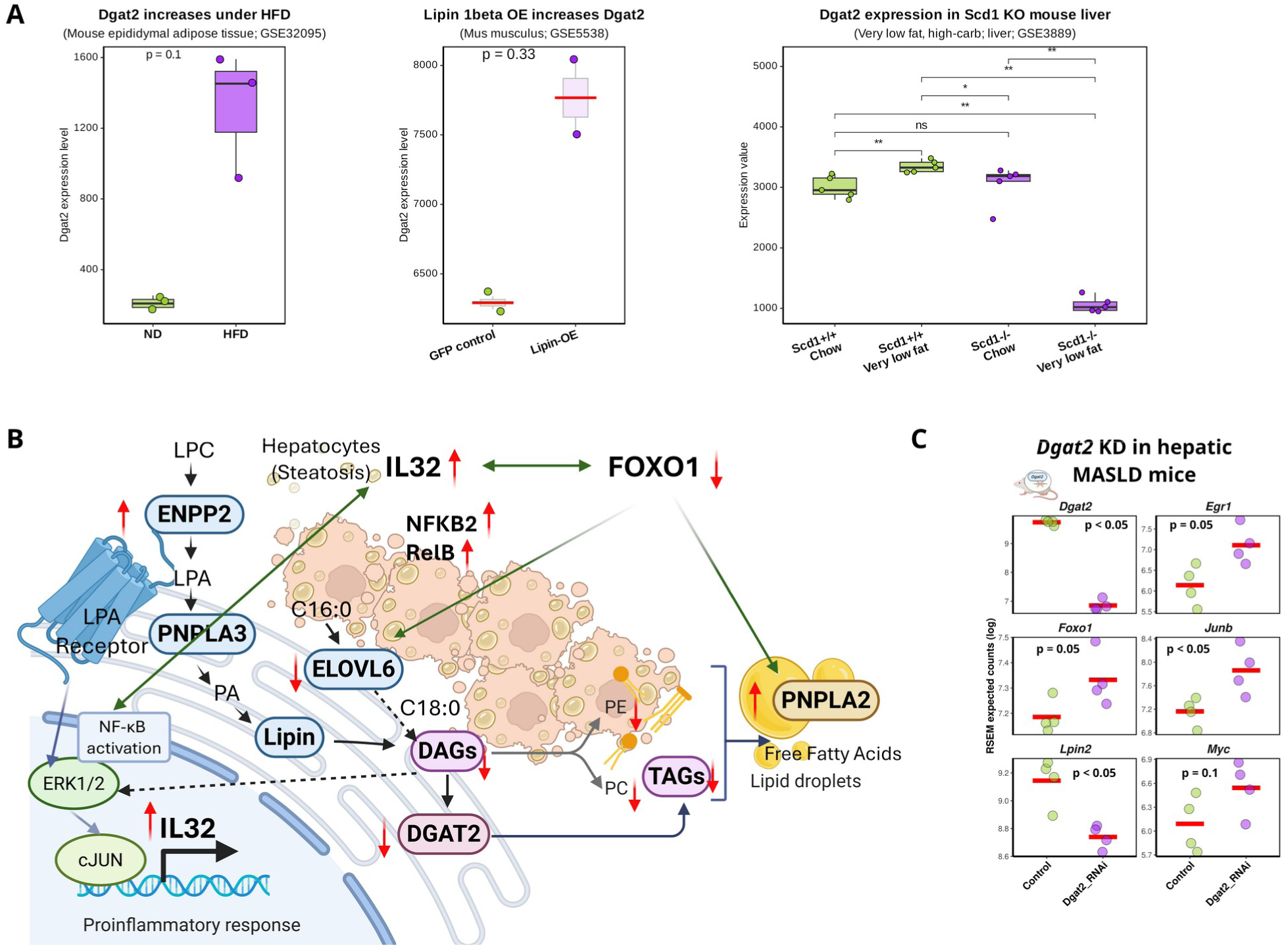
Metabolic reprogramming of TAG synthesis in HCC. (A) Dgat2 expression is induced by high-fat diet (GSE32095) and Lipin 1-beta overexpression (GSE5538) but suppressed by Scd1 deficiency (GSE3889) in mouse liver and adipose tissue. Red bars indicate the mean. Statistical comparisons as shown. (B) Schematic overview of altered metabolomic fluxes in HCC tumors versus non-tumor liver from MASLD-HCC patients. Illustration created with BioRender. (C) Differential expression of metabolic and NF-κB/ERK-related genes following *Dgat2* knockdown in genetically obese MASLD mice treated with RNAi targeting hepatic *DGAT2*.

Consistent with these findings, human HCC samples exhibit significant DGAT2 downregulation at both the transcript and protein levels in tumors compared to adjacent non-tumoral liver tissue, accompanied by a marked reduction in TAGs, DAG substrates, and downstream phospholipid species including PC and PE (**Fig. 2A-B, 2F**).

Genetic ablation of DGAT2 in Huh7 cells primarily affects pathways linked to the DAG-TAG flux, including reduced expression of LPCAT1 and LPIN2 genes similarly to our observations in mice (**Fig. 6C, S5a**). Notably, DGAT2 loss alone is insufficient to suppress FAO or recapitulate the broad depletion of storage and structural lipids observed in tumors (**Fig. 2E-F**, **S2b**), indicating that DGAT2 is not alone in driving this global metabolic remodeling.

At the cell-autonomous level, DGAT2 knockout Huh7 cells display significant rewiring, rather than uniform activation, of NF-κB and ERK transcriptional programs, including downregulation of MYC, EGR1, and ETVF, and upregulation of CXCL10, changes that contrast with the activation observed in murine MASLD but reflect a context-dependent adjustment of signaling outputs (**Fig. S5a**).

In summary, DGAT2 loss alone is insufficient to drive global metabolic remodeling or full NF-κB/ERK activation, but instead reprograms lipid metabolism to create a permissive state that sensitizes hepatocytes to IL-32-driven inflammatory signaling.

## DISCUSSION

A central finding of this study is that IL-32 transcription is governed by a switch from canonical to non-canonical NF-κB signaling. The NF-κB pathway is classically activated in HCC, functioning as a central adaptive transcriptional program that integrates tumor progression, metabolic rewiring, and immune evasion (18). While canonical NF-κB (RELA/NFKB1) has traditionally been considered the primary driver of inflammatory gene expression in HCC, we identify NFKB2/RELB, which is less defined than canonical NF-κB in HCC (22), as the dominant regulatory module, forming a self-reinforcing inflammatory circuit that sustains IL-32 expression. This is consistent with emerging evidence that RELB drives pro-metastatic programs in the liver TME (37), but critically, we defined that this non-canonical axis is the primary driver of IL-32 in HCC, whereas canonical signaling plays only a permissive basal role.

This NF-κB module switch is not simply transcription factor substitution but reflects a broader remodeling of chromatin accessibility. We show that FOXO1 functions as a pioneer factor that maintains open chromatin at *IL32* and NF-κB regulatory loci, thereby licensing the inflammatory switch. Recent studies have demonstrated that FOXO1 binds condensed chromatin regions and induces local epigenetic remodeling to promote memory programs and restrain exhaustion (38). Here, we extend this paradigm and demonstrate that FOXO1 acts as a pioneer factor for the non-canonical NF-κB inflammatory switch in HCC, linking metabolic state to inflammatory competence. Loss of FOXO1 disrupts this accessibility landscape, positioning FOXO1 as a key epigenetic gatekeeper of IL-32 signaling.

Beyond transcriptional regulation, IL-32 is embedded in a lipid-inflammatory interface centered on DGAT2. Although DGAT2 expression strongly correlates with IL-32 and reflects hepatic lipid storage capacity, its loss is insufficient to recapitulate global metabolic remodeling in HCC. Instead, DGAT2 depletion rewires NF-κB and ERK signaling in a context-dependent manner, suggesting that DGAT2 functions primarily as a permissive node that modulates cellular sensitivity to inflammatory cues rather than acting as a primary driver of metabolic collapse. This aligns with emerging evidence that DGAT2 sustains mitochondrial stability via transcriptional network and suppresses glycolytic metabolic shifts in HCC (39).

Importantly, IL-32 regulation is not restricted to inflammatory signaling but integrates metabolic inputs, as evidenced by its responsiveness to PPAR signaling and lipid state-dependent transcriptional programs. IL-32 has been shown to enhance lipid accumulation and inhibit cholesterol efflux in macrophages via the PPARγ-LXRα-ABCA1 pathway (40), establishing a bidirectional metabolic-inflammatory interface.

Collectively, our findings define IL-32 as a context-dependent regulatory hub that integrates metabolic state and inflammatory signaling during hepatocarcinogenesis. We propose a model in which FOXO1-dependent chromatin priming enables a non-canonical NF-κB-driven inflammatory switch, while lipid metabolic rewiring via DGAT2 loss shapes the sensitivity and output of this circuit. This coordinated reprogramming shifts IL-32 from a metabolic regulator in MASLD to an inflammation-amplifying node in HCC, a functional transition that has not been previously appreciated.

### Limitations of the current study

Two mechanistic questions remain. First, our FOXO1 chromatin-priming and NF-κB switch studies were conducted primarily in macrophage and hepatoma cell lines; whether FOXO1 binding instructs NFKB2 recruitment or stabilizes pre-existing accessible states, and whether this pioneer activity is equally robust across primary human hepatocytes and other hepatic immune subsets, requires temporal and cell-type-specific resolution. Second, although DGAT2 acts as a permissive inflammatory node, the specific bioactive lipid species mediating its context-dependent NF-κB and ERK rewiring, and whether this function is shared across hepatic compartments, remain undefined. Despite these nuances, our integrated framework positions the FOXO1-NF-κB-IL-32 axis as a therapeutic link between metabolic state and inflammatory competence in MASLD-HCC.

### Author contributions

Conceptualization: L.N.Z, and J.B.A; Data acquisition: L.N.Z, P.K. and J.B.A. Data analysis and visualization: L.N.Z; Writing: L.N.Z, P.K and J.B.A. Funding: P.K. and J.B.A.

#### Data availability

The RNA-seq for MASLD are available through GEO under accession number GSE281797. The untargeted LC-HRMS data for MASLD have been deposited in Zenodo (DOI 10.5281/zenodo.14091962), SWATH data for the organic fraction (DOI 10.5281/zenodo.14096635), and SWATH data for the aqueous fraction (DOIs 10.5281/zenodo.14096753 and 10.5281/zenodo.14136832). All clinical data, processed omics datasets, and relevant code related to MASLD cohort are accessible at https://github.com/SLINGhub/MASLD_dual_omics. The RNA-seq for MASLD-associated HCC has been deposited at GEO under accession number GSE334651.

## ACKNOWLEDGMENTS

The authors would like to thank the patients and their families for allowing research access to their samples. Also, we are grateful for access to all public data sets. The results are in part supported by data generated by the TCGA Research Network (https://www.cancer.gov/tcga), Gene Expression Omnibus (GEO), European Nucleotide Archive (ENA), and the Encyclopedia of DNA Elements (ENCODE) . The authors would like to express their gratitude to Profs. Hyungwon Choi (Metabolic data), Matthew Watt (sample collections), and Patrik Midlöv for support and help during the study.

## Supplementary Figure Legends

Figure S1. *IL32* expression is elevated in MASLD and correlates with hepatic TAG remodeling.

Figure S2. Lipid and metabolic gene alterations in MASLD-HCC tumors.

Figure S3. Effects of *NFKB1* loss on NF-κB signaling and related pathways.

Figure S4. Mechanisms of FoxO1 network regulation and HCC-associated suppression.

Figure S5. Transcriptomic analysis of *DGAT2*-deficient hepatocytes and *IL32*-related transcriptional networks.

