## Supplemental text and figures for "IL-32 is a stress-responsive node locked in a noncanonical NF-κB inflammatory loop during MASLD to HCC transition": supportingMaterial.IL32_v9.docx

Supporting Material

**METHOD**

### **Study design and clinical cohorts.** Full details of cohort recruitment, inclusion/exclusion criteria, liver biopsy collection, histological assessment, and metabolomics sample preparation have been described previously(1). In brief, eligible patients with obesity scheduled for bariatric surgery (sleeve gastrectomy, gastric bypass, or laparoscopic‑adjustable band placement) were prospectively enrolled. Exclusion criteria included other causes of chronic liver disease, significant alcohol intake (AUDIT score), recent use of steatogenic medications, and liver biopsy weight <2 mg for omics analyses. Two in‑house clinical cohorts were analyzed: a MASLD cohort (steatosis grade ≥1 with or without lobular inflammation) and a MASLD‑associated HCC cohort. Venous blood was collected after an overnight fast for clinical biochemistry and plasma metabolomics. Wedge liver biopsies (~1 cm³) were obtained from the left lobe during surgery; one portion was formalin‑fixed for histology (NAS score and Kleiner fibrosis staging), and the remainder was snap‑frozen for multi‑omics analysis.

### **Untargeted metabolomics.** For liver tissue (50 ± 5 mg) and plasma (50 µL), metabolites and lipids were extracted using a methanol:chloroform:water phase separation protocol as previously described (1). Briefly, liver samples were homogenized with methanol:chloroform (50:50 v/v), phase separation was induced by adding water, and the upper (polar metabolites) and lower (lipids) phases were dried under SpeedVac. Plasma samples were extracted with methanol and chloroform, followed by water addition and centrifugation. Dried extracts were reconstituted in 90:10 isopropanol:acetonitrile (organic fraction) or 90% acetonitrile (aqueous fraction), sonicated, and centrifuged before LC‑MS analysis.

**Single cell RNA-seq processing and analysis.** Publicly available single cell RNA-seq datasets obtained from NASH-related HCC **(GSE189175)(2)**. scRNA-seq datasets of primary and metastatic HCC (GSE149614 and GSE189903)(3,4). **The analysis was performed on paired human HCC and non-tumor liver samples. Raw data were processed, doublets were removed using scDblFinder and were normalized and integrated via the SCTransform workflow.**

**Spatial transcriptomic analysis of MASLD liver tissue.** Processed spatial transcriptomic data from two MASLD liver slides were independently analyzed using standard Seurat workflows (GSE292268)(5).

**Public genomic and transcriptomic data.** To investigate the transcriptional mechanisms underlying our metabolomic observations, we analyzed relevant public datasets. This included RNA-Seq and ChIP-Seq data from FOXO1-overexpressed HepG2 cells (GSE255416)(6), RNA-Seq data from NFKB1-knockdown MCF7 cells(7), and transcriptomic datasets related to DGAT2 manipulation and inflammasome activation (GSE163723(8), GSE178987(9)). To capture disease-relevant inflammatory stimuli, we additionally incorporated transcriptomic profiles from alcoholic hepatitis (GSE28619(10)), hepatitis C virus core protein stimulation (GSE2067(11)), and TGF-β–treated hepatocytes (GSE10393(12)), which represent progressive inflammatory and fibrotic states of liver injury.

To assess metabolic regulatory inputs, we integrated datasets probing hepatocyte identity and lipid-sensing pathways, including HNF4A RNAi knockdown in HepG2 cells (GSE29084(13)), PPAR ligand-response profiling (GSE2699(14)), lipin1β overexpression in mouse liver (GSE5538(15)), and Scd1 knockout mice under low-fat dietary conditions (GSE3889(16)). In addition, high-fat diet–associated transcriptional remodeling was incorporated through DGAT2- and lipid metabolism-linked datasets, including GPR120-mediated responses to high-fat diet (GSE32095(17)).

To provide regulatory context, we incorporated **ChIP-Seq** data of **RELA (ENCFF335FKX), REL (ENCFF367AVM), NFKB2 (ENCFF918FFX), HNF4A** from HepG2 cells, along with **epigenomic signals** from HepG2 **ATAC-Seq** (ENCFF285FQS) and **CAGE** (ENCFF205BRW) datasets, sourced from ENCODE. Finally, we incorporated lipidomic data from *IL-32* CRISPR/Cas9 knockout versus wild-type cells(18) to link transcriptional regulation with metabolic outcomes.

**Analysis of FOXO1 knockout in human HepG2 cells and **murine model**.** Public RNA-seq data from FOXO1-knockout HepG2 cells stimulated with lipopolysaccharide (LPS) and corresponding wild-type controls were obtained from the GEO (GSE293003)(19). Ensembl gene identifiers were mapped to official HGNC gene symbols using the biomaRt package(20).

To validate findings in vivo, we used transcriptomic data from a Foxo1-knockout mouse model (GSE90754). A similar analytical workflow was applied. Mouse Ensembl identifiers were converted to standard gene symbols via biomaRt. Processed expression values were structured for comparative analysis between knockout and control groups.

**NFKB1 knockout in mice.** Microarray data (CEL files) from an Nfkb1 knockout mouse model were obtained from the Gene Expression Omnibus (GSE23497)(21). Raw data were normalized using the Robust Multi-array Average (RMA) method.

**NFKB2 overexpression analysis in mice.** Public RNA-seq raw count data from a murine model of NFKB2 overexpression versus control were obtained from the GEO (GSE250065)(22). Differential expression analysis was performed using the DESeq2 package(23) in R. The DESeq2 workflow was executed using default parameters to estimate size factors, dispersions, and fit negative binomial generalized linear models. For downstream interpretation, the Ensembl IDs of the top 100 differentially expressed genes ranked by absolute log2 fold change were converted to mouse gene symbols using the biomaRt package(24). Pathway enrichment analysis of this gene list was subsequently performed using Metascape(25).

***Dgat2* knockout in MASLD mouse model.** RNA-seq data obtained from the *Dgat2* knockout model of MASLD (ENA: ERP114849) was processed to define experimental conditions based on genotype and treatment. A DESeq2 object was created with the appropriate design, and the DESeq2 workflow was executed to perform differential expression analysis between conditions. Validation was performed using processed RNA-seq data from **Dgat2 RNAi knockdown in mouse liver** (GSE178987)(26).

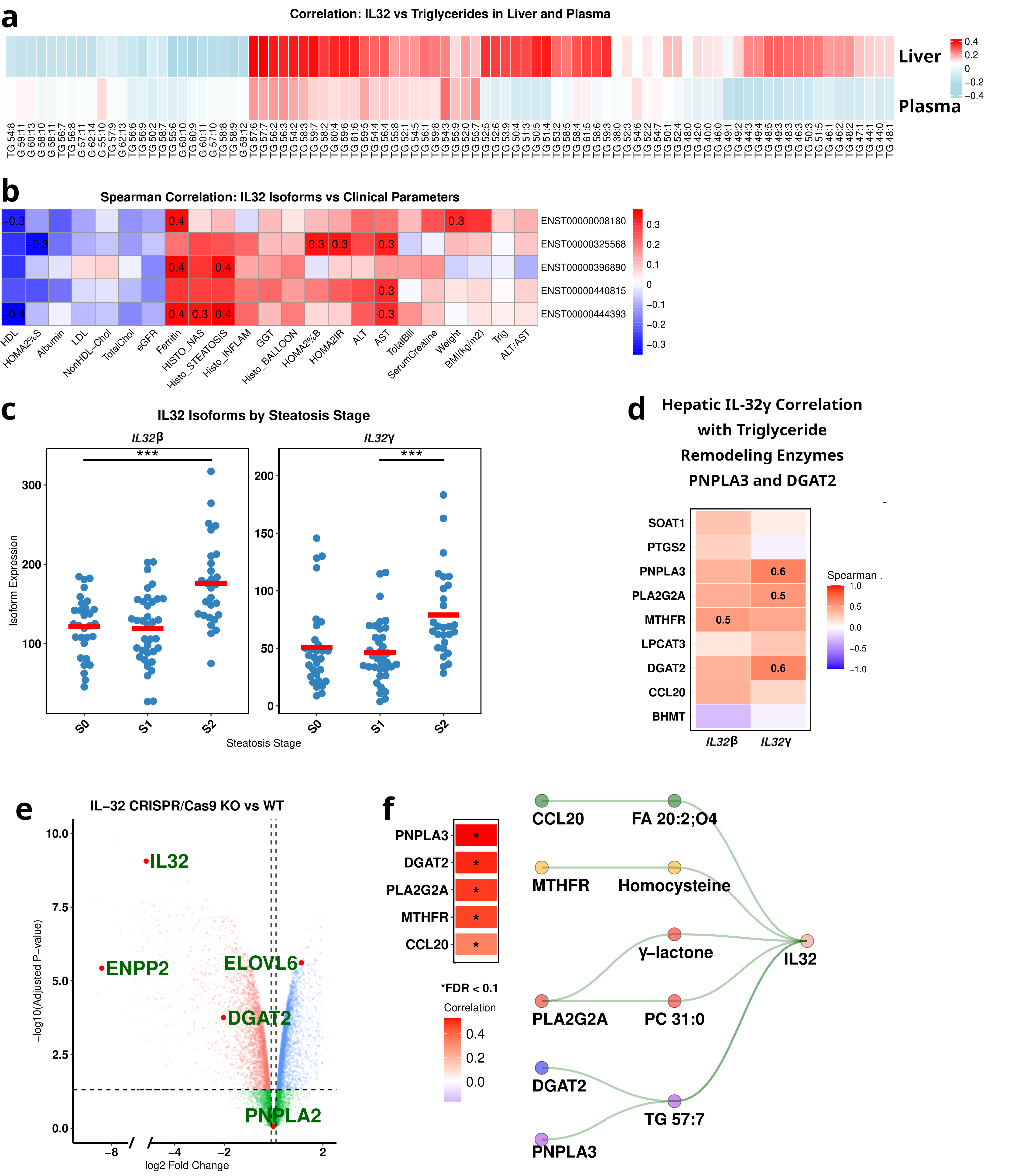

Fig. S1. **IL-32 expression is elevated in MASLD and correlates with hepatic triglyceride remodeling.**

**(a)** Correlation analysis of hepatic and plasma IL-32 levels with intrahepatic and circulating TAG species in the MASLD cohort.

**(b)** Association of hepatic IL-32 expression with clinical parameters in individuals with MASLD.

**(c)** IL-32β and IL-32γ isoform expression increases with steatosis severity in human MASLD liver.

**(d)** IL-32γ shows significant positive correlation with expression of the TAG remodeling enzymes DGAT2 and PNPLA3 in MASLD.

(e) Volcano plot showing differential metabolic genes between IL-32 CRISPR/Cas9 KO and WT cells.

(f) Significant t**op correlations between hepatic IL-32 expression and** IL-32-centered network in MASLD.

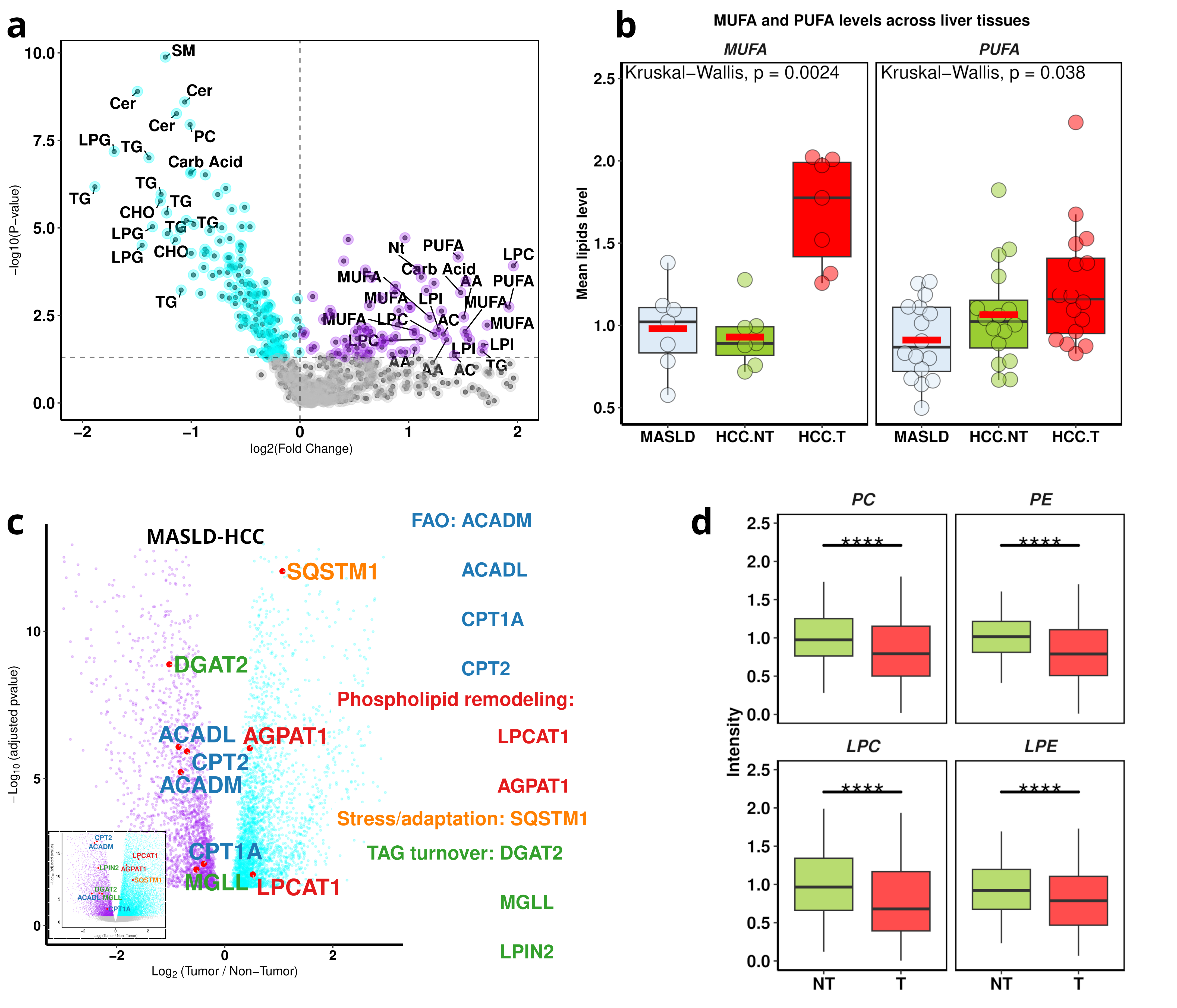

**Fig. S2. Lipid and metabolic gene alterations in MASLD-HCC tumors.**

**(a) Volcano plot showing downregulated and upregulated metabolites in HCC tumors versus non-tumor tissue (HCC.T vs. HCC.NT). The x-axis shows the log₂ fold change (log₂FC) and the y-axis shows the −log₁₀(p-value).**

**(b)** Increased levels of **PUFAs and MUFAs** in hepatocellular carcinoma tumors (MASLD-HCC).

(c) Differential expression of key metabolic genes in MASLD-HCC tumor versus non-tumor tissue (Bottom left panel: TCGA-LIHC). Volcano plot showing log₂ fold changes versus -log₁₀ adjusted p-values for all genes. Metabolic genes of interest are highlighted and labeled, with colors indicating functional category: **FAO** (blue), **TAG turnover** (green), **Phospholipid remodeling** (red), and **Stress/adaptation** (orange). This visualization highlights the coordinated downregulation of fatty acid oxidation and triglyceride metabolism genes, alongside upregulation of phospholipid remodeling and stress-response genes in tumors.

**(d)** Tumor hepatocytes exhibit a **coordinated downregulation of phospholipids**, including **PC, PE, LPC, and LPE** (MASLD-HCC).

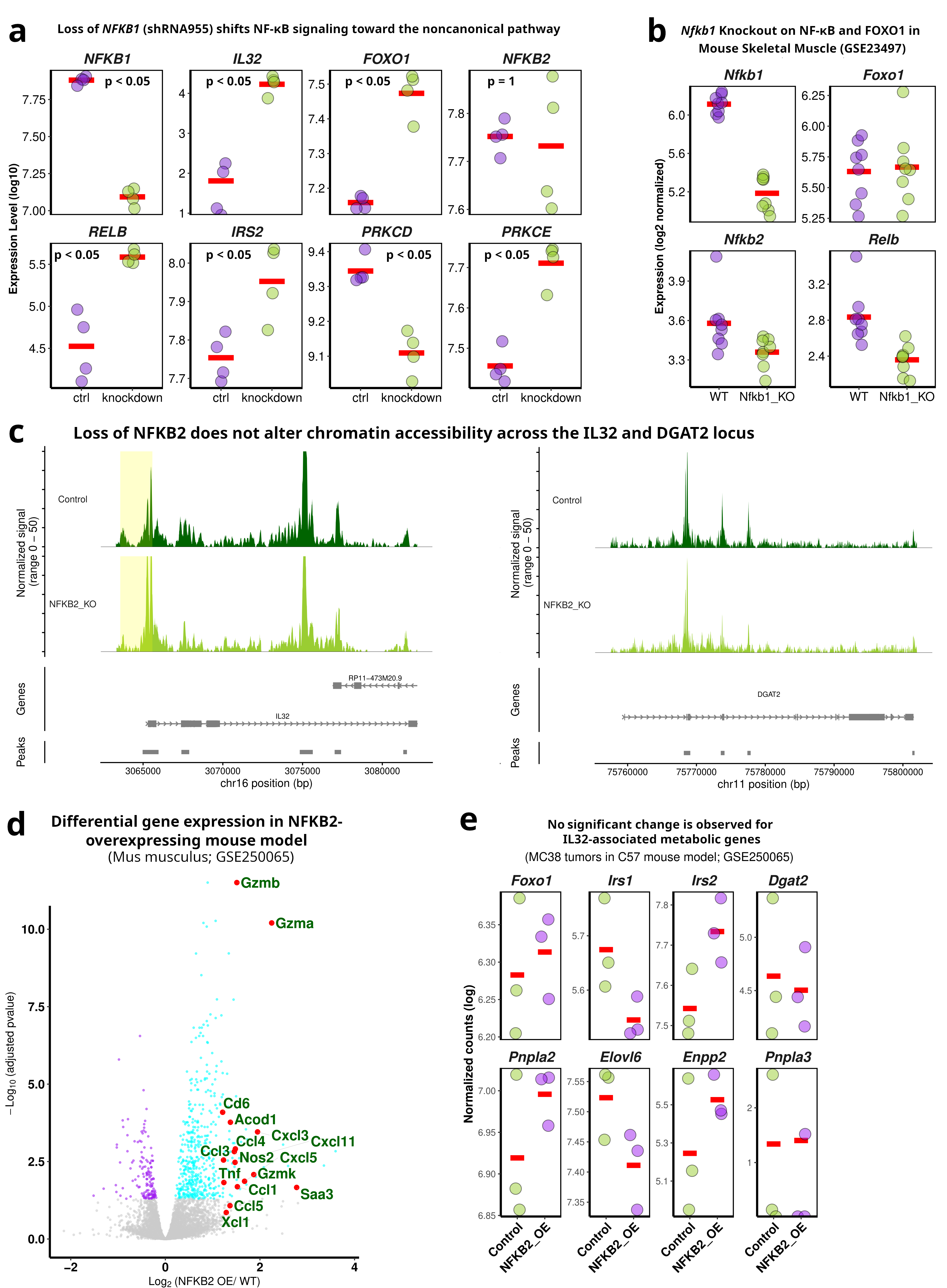

**Fig. S3. Effects of NFKB1 loss on NF-κB signaling and related pathways.**

**(a)** Loss of NFKB1 redirects NF-κB signaling toward the noncanonical pathway. Log-transformed expression levels of key inflammatory, metabolic, and signaling genes in control versus NFKB1 knockdown (shRNA955; shRNA954, see Fig. 2E) in MCF7 cells.

**(b)** Effects of Nfkb1 knockout on NF-κB and Foxo1 expression in mouse skeletal muscle (GSE23497).

(c) Plot showing chromatin accessibility across the IL32 and DGAT2 loci in NFKB2-knockout versus non-targeting control T cells, generated from ASAP-seq pooled CRISPR perturbation data (GSE156476). Tracks display ATAC-seq signal smoothed with a 100-bp window and extended 2kb upstream and 0.5kb downstream of each gene, with MACS2 peaks shown above each profile, and cells grouped by genotype. The accessibility profiles indicate that loss of **NFKB2** does not produce detectable changes in chromatin accessibility at the IL32 promoter or across the broader IL32 locus.

(d) Volcano plot showing differentially expressed genes between NFKB2-overexpressing and wild-type cells. Genes associated with inflammation are highlighted (GSE250065).

(e) Expression of IL32-associated metabolic genes in MC38 tumors from C57 mice with or without NFKB2 overexpression (GSE250065). Gene expression was measured as normalized counts (log-transformed). No significant changes were observed in the expression of these metabolically linked genes upon NFKB2 overexpression, indicating that NFKB2 primarily influences inflammatory and immune pathways rather than metabolic gene expression.

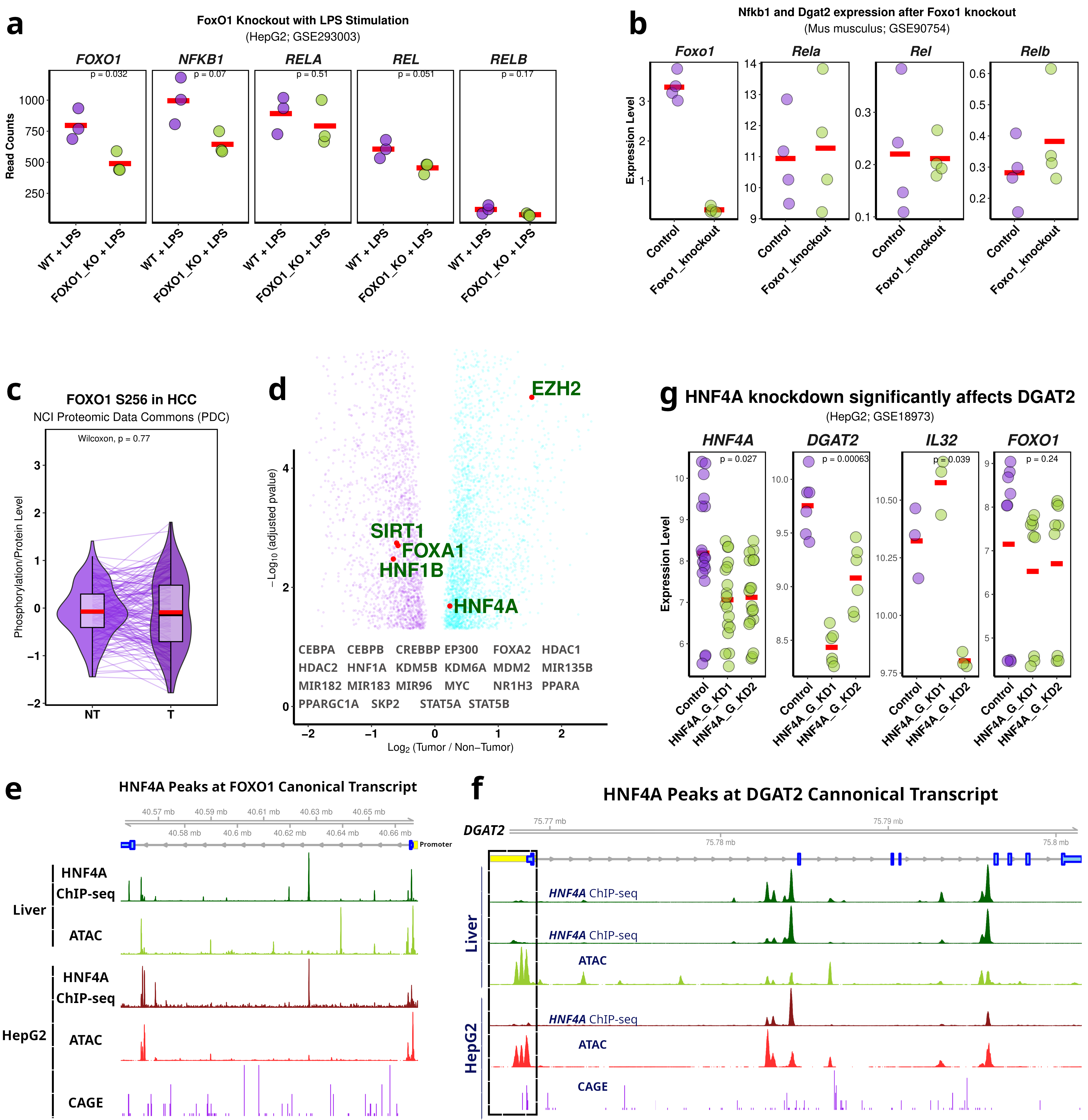

Fig. S4. **Mechanisms of FoxO1 network regulation and HCC-associated suppression.**

(a) *FoxO1* knockout in LPS-stimulated human HepG2 hepatocytes (GSE293003) confirms a broad suppression of the NF-κB pathway (NFKB1, RELA, REL, RELB). Related to Fig. 4C.

(b) *FoxO1* knockout in vivo. Related to Fig. 4D

(c) **FOXO1 phosphorylation at Ser256 remains stable in hepatocellular carcinoma.**

**(d) Altered FOXO1 regulators in MASLD-HCC. Volcano plot identifies transcription factors and chromatin modifiers linked to FOXO1 control that are differentially expressed in HCC tumors.**

(e) Genomic landscape of the FOXO1 locus with liver regulatory features. Genome browser visualization showing the FOXO1 gene track alongside HNF4A ChIP-seq and ATAC-seq profiles from healthy liver, highlighting transcription factor binding and chromatin accessibility at the locus. Corresponding HNF4A-associated ATAC-seq and CAGE signals from HepG2 cells demonstrate conserved regulatory activity in human liver-derived cancer cells. The FOXO1 promoter region is indicated by a yellow-highlighted box marking the primary transcription start site and its overlap with HNF4A-linked accessible chromatin.

(f) Genomic landscape of the DGAT2 locus with liver regulatory features.

(g) Expression profile of FOXO1, DGAT2, and IL32 following HNF4A knockdown (HepG2; GSE18973).

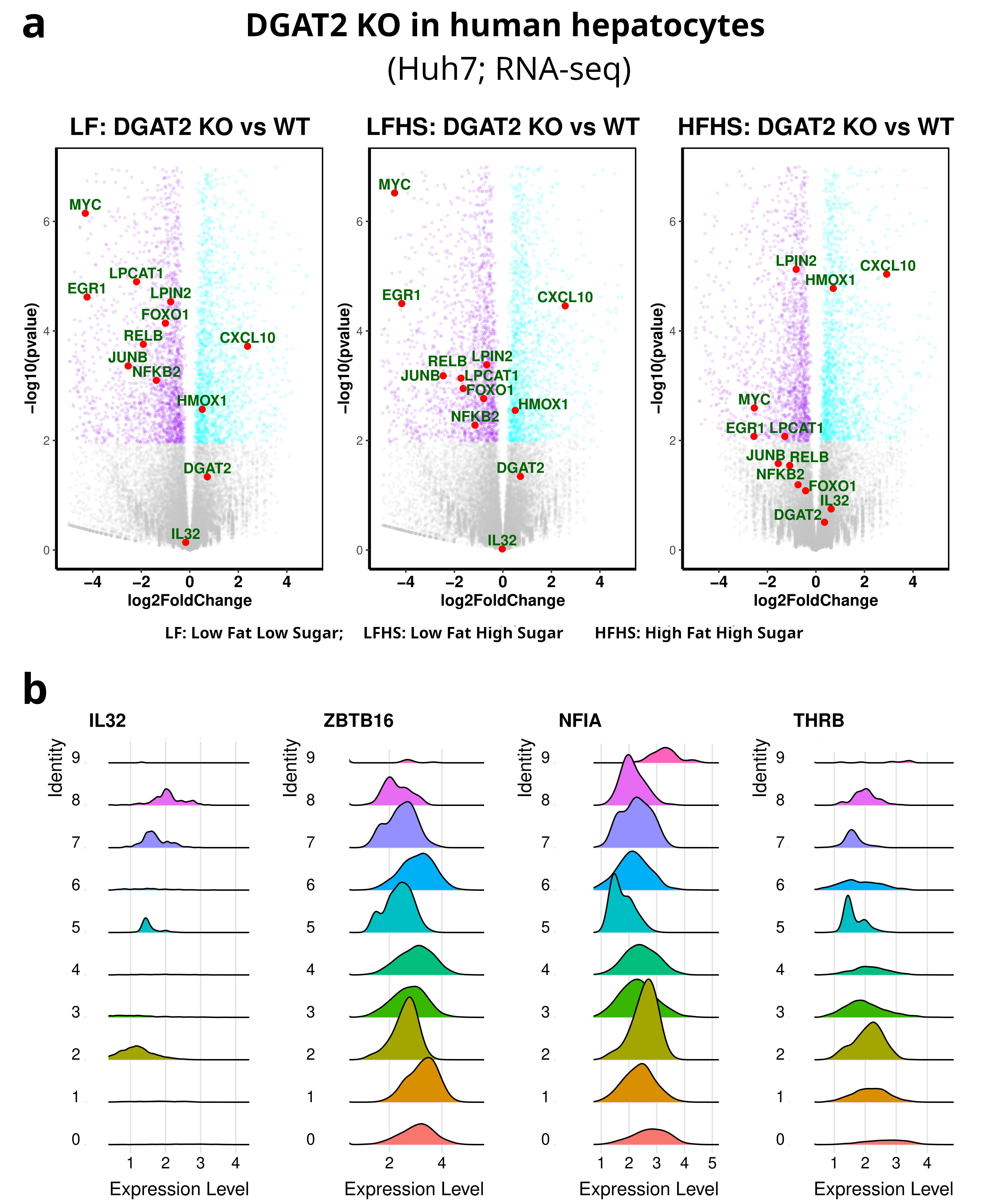

Fig. S5. Transcriptomic analysis of DGAT2-deficient hepatocytes and IL32-related transcriptional networks.

(a) Differential expression of metabolic and NF-κB/ERK-related genes in DGAT2 knockout human hepatocytes (Huh7) under low fat low sugar (LF), low fat high sugar (LFHS), and high fat high sugar (HFHS) conditions, compared with wild-type cells. RNA-seq data from ENA: ERP114849.

(b) **Ridge plots showing single-nucleus RNA-seq expression of IL32 and candidate nuclear binding partners.** Expression of IL32 and the transcription factors ZBTB16, NFIA, and THRB across cell populations in the dataset.
